# Integrated Analysis of Gene Expression Differences in Twins Discordant for Disease and Binary Phenotypes

**DOI:** 10.1101/226449

**Authors:** Sivateja Tangirala, Chirag J Patel

## Abstract

While both genes and environment contribute to phenotype, deciphering environmental contributions to phenotype is a challenge. Furthermore, elucidating how different phenotypes may share similar environmental etiologies also is challenging. One way to identify environmental influences is through a discordant monozygotic (MZ) twin study design. Here, we assessed differential gene expression in MZ discordant twin pairs (affected vs. non-affected) for seven phenotypes, including chronic fatigue syndrome, obesity, ulcerative colitis, major depressive disorder, intermittent allergic rhinitis, physical activity, and intelligence quotient, comparing the spectrum of genes differentially expressed across seven phenotypes individually. Second, we performed meta-analysis for each gene to identify commonalities and differences in gene expression signatures between the seven phenotypes. In our integrative analyses, we found that there may be a common gene expression signature (with small effect sizes) across the phenotypes; however, differences between phenotypes with respect to differentially expressed genes were more prominently featured. Therefore, defining common environmentally induced pathways in phenotypes remains elusive. We make our work accessible by providing a new database (*DiscTwinExprDB*: http://apps.chiragjpgroup.org/disctwinexprdb/) for investigators to study non-genotypic influence on gene expression.

## Introduction

Gene expression is influenced by both inherited and non-inherited (or environmental) factors; however identifying how environment influences phenotype, such as disease, remains a challenge^1^. A common approach to identify differentially expressed genes in disease is the case-control study. Case-control studies involve the matching of affected individuals with healthy controls to assess the differences of gene expression in cases versus controls. However, it is difficult to identify the causes of differences of gene expression with respect to inherited factors, environmental, or phenotypic state; further, associations may be biased due to confounding variables.

One way to partition the role of environment and inherited factors in gene expression is to use a family-based twin-design, whereby twins are discordant for phenotypes. For example, monozygotic (MZ) discordant twins are twins that share the same genome but are discordant for a phenotype (e.g., one twin has a certain phenotype, the other does not). The monozygotic discordant twin study design provides a natural study design in order to identify significant genes for a particular phenotype after controlling for non-temporally dependent variables, such as shared genetics, sex, and age^2^.

Is there a consistent gene expression signature of environmental influence? Or, how much does gene expression due to potential environmental influence vary across phenotypes? We hypothesized that integrating gene expression data from multiple phenotypes can allow the elucidation of heterogeneity of discordant gene expression (how gene expression differences between twins vary) and furthermore, gene signatures across phenotypes. More specifically, we claim it is possible to measure cross-phenotype heterogeneity by meta-analyzing across mean expression differences for each gene from discordant twin samples. As of this writing, gene expression data from discordant twin samples have not been utilized to perform such analyses.

Our study’s goal was to identify significant differentially expressed genes between samples of affected and non-affected MZ twin pairs and integrate mean expression differences across seven phenotypes. We hypothesize that it is possible to detect potential environmentally modulated gene expression values shared and distinct among different phenotypes. We further claim that identifying genes that are different and shared among numerous phenotypes will shed light on shared environmental etiology in phenotypic variation.

In order to identify genes in discordant twins, we formulated a computational approach in four parts. First, we queried public repositories such as the Gene Expression Omnibus^4^ (GEO) for gene expression studies of discordant monozygotic twin pairs. Next, we identified significantly altered genes in twin pairs that are discordant for each of the seven phenotypes. We then compared gene signatures across the seven phenotypes in a pairwise fashion and by using a meta-analytic approach.Last, we hypothesized that sex may also play a role in gene expression variation; thus, we attempted to identify genes in a sex-specific manner across multiple phenotypes.

## Results

### Differential gene expression analysis in each of the seven phenotypes individually

For our gene expression analyses we used expression data from the Gene Expression Omnibus^4^ (GEO), Array Express^5^ (AE), and a study from the Database of Genotypes and Phenotypes^6^ (dbGAP) study (Fig. 1, Table 1). The seven phenotypes we interrogated included 4 diseases such as chronic fatigue (CFS), major depressive disorder (MDD), ulcerative colitis (UC), intermittent allergic rhinitis (IAR in vitro), and 3 phenotypes including physical activity (PA), obesity (OB), intelligence quotient (IQ). To ensure adequate power for detection, we used seven studies that each had at least 10 twin pair samples. The sample sources (tissues and cell lines) used by the studies included peripheral blood, lymphoblastoid cell lines, adipose tissue, muscle tissue, and colon tissue (Table 1). All results are accessible via our R Shiny web application (http://apps.chiragjpgroup.org/disctwinexprdb/).

**Figure 1.**
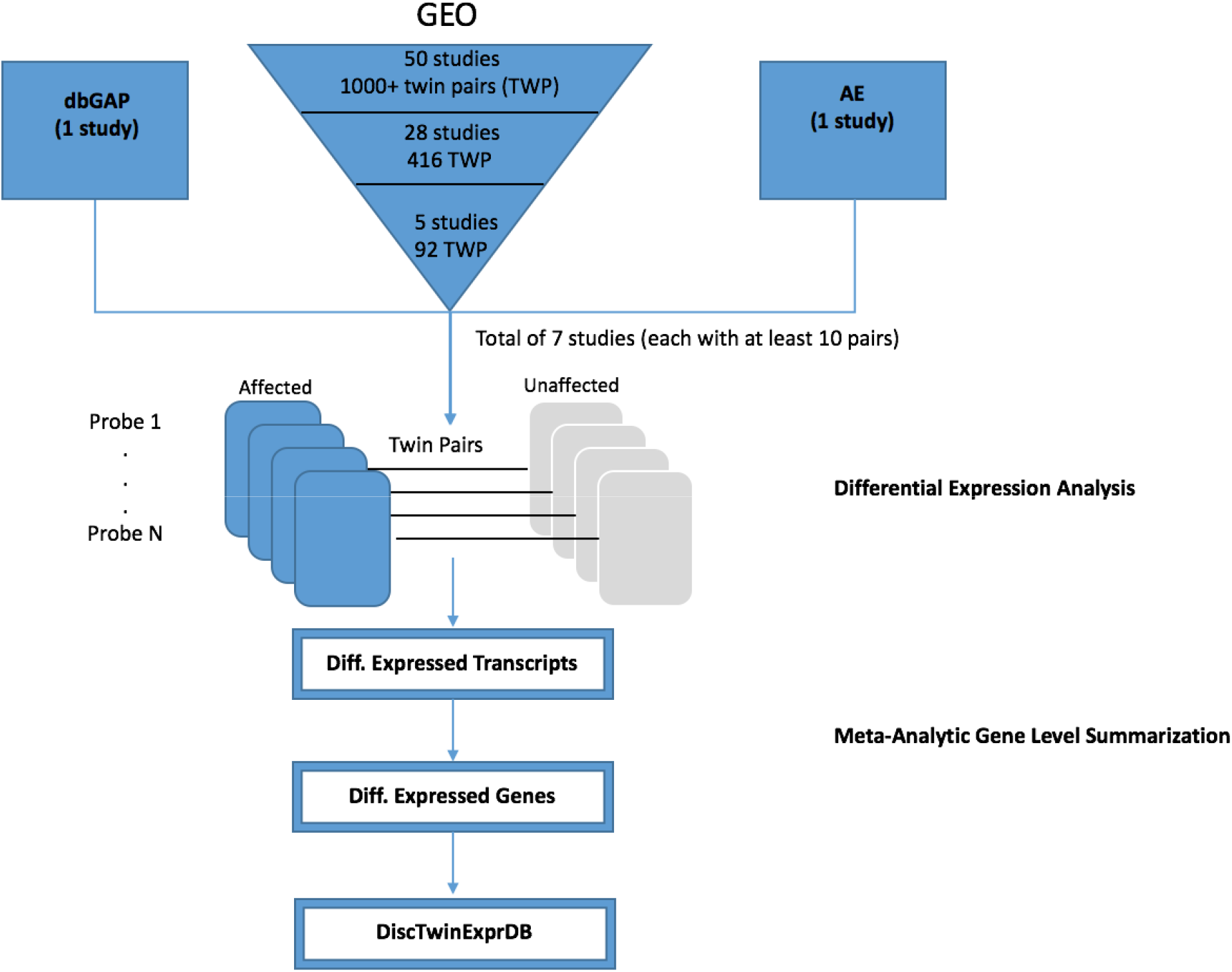
Analysis Procedure. A schematic diagram depicting the analysis pipeline.(1) Data Selection involved a filtration process for selecting twin expression datasets (2) Differential Expression Analysis was carried out (using probe or transcript-level values) to find significant differentially expressed transcripts using FDR and effect size thresholds (3) Meta-Analytic Gene Level Summarization was carried out to summarize transcript-level differences to gene-level differences

**Table 1:** Summary of Datasets. This table shows the phenotype and number of genes being measured, sample size, platform, tissue, source, and reference paper for each of the seven studies.

| Study identifier | Reference (s) | Number of Twin Pairs | Phenotype | Number of Genes | Platform | Sample Source( Tissue and cell lines) | Source |
| --- | --- | --- | --- | --- | --- | --- | --- |
| GSE22619 | Lepage *et al.*<sup>21</sup>,<br>Häsler *et al.*<sup>22</sup> | 10 | ulcerative colitis (UC) | 22836 | GPL570 | Primary mucosal tissue, colon | GEO |
| GSE16059 | Byrnes *et al.*<sup>9</sup> | 44 | chronic fatigue syndrome (CFS) | 22836 | GPL570 | Peripheral venous blood | GEO |
| GSE20319 | Leskinen *et al.*<sup>23</sup> | 10 | physical activity (PA) | 19429 | GPL6884 | Musculus vastus lateralis | GEO |
| GSE33476 | Yu *et al.*<sup>24</sup> | 17 | intelligence quotient(IQ) | 18638 | GPL6244 | Lymphoblastoid cell lines | GEO |
| GSE37146 | Sjogren *et al.*<sup>25</sup> | 11 | intermittent allergic rhinitis (in vitro) (IAR_invitro) | 19580 | GPL6102 | Peripheral blood mononuclear cells | GEO |
| MDD(dbGAP) | Wright *et al.*<sup>26</sup> | 28 | major depressive disorder (MDD) | 19284 | GPL13667 | Peripheral blood | dbGAP |
| E-MEXP-1425 | Pietiläinen *et al.*<sup>27</sup> | 13 | obesity (OB) | 22836 | GPL570 | Adipose tissue | Array Express |

We performed differential gene expression analysis on each of the monozygotic discordant twin gene expression studies, performed meta-analysis of probe-level (transcript-level) values to obtain gene-level values^7^, and corrected the p-values for each gene-level value using the false discovery rate (FDR) method^15^. In order to minimize the false positive rate (FPR), Sweeney et al. suggested to utilize stringent significance and effect size thresholds^7^. Therefore, we identified significant differentially expressed genes for each phenotype that fell under a FDR threshold of 0.05 and had an effect size threshold greater than the 95th percentile of the absolute value of mean gene expression differences in each phenotype (Fig. 2, Fig. S1). Figure S1 shows the empirical cumulative distribution of mean differences for each phenotype. The number of significant genes ranged from a total of three significant in chronic fatigue syndrome (CFS) to 677 in intelligence quotient (IQ). Overall, the total number of unique significant genes across all the seven datasets was 1,286 out of the 25,154 total number of genes measured across all of those datasets (5%).

**Figure 2.**
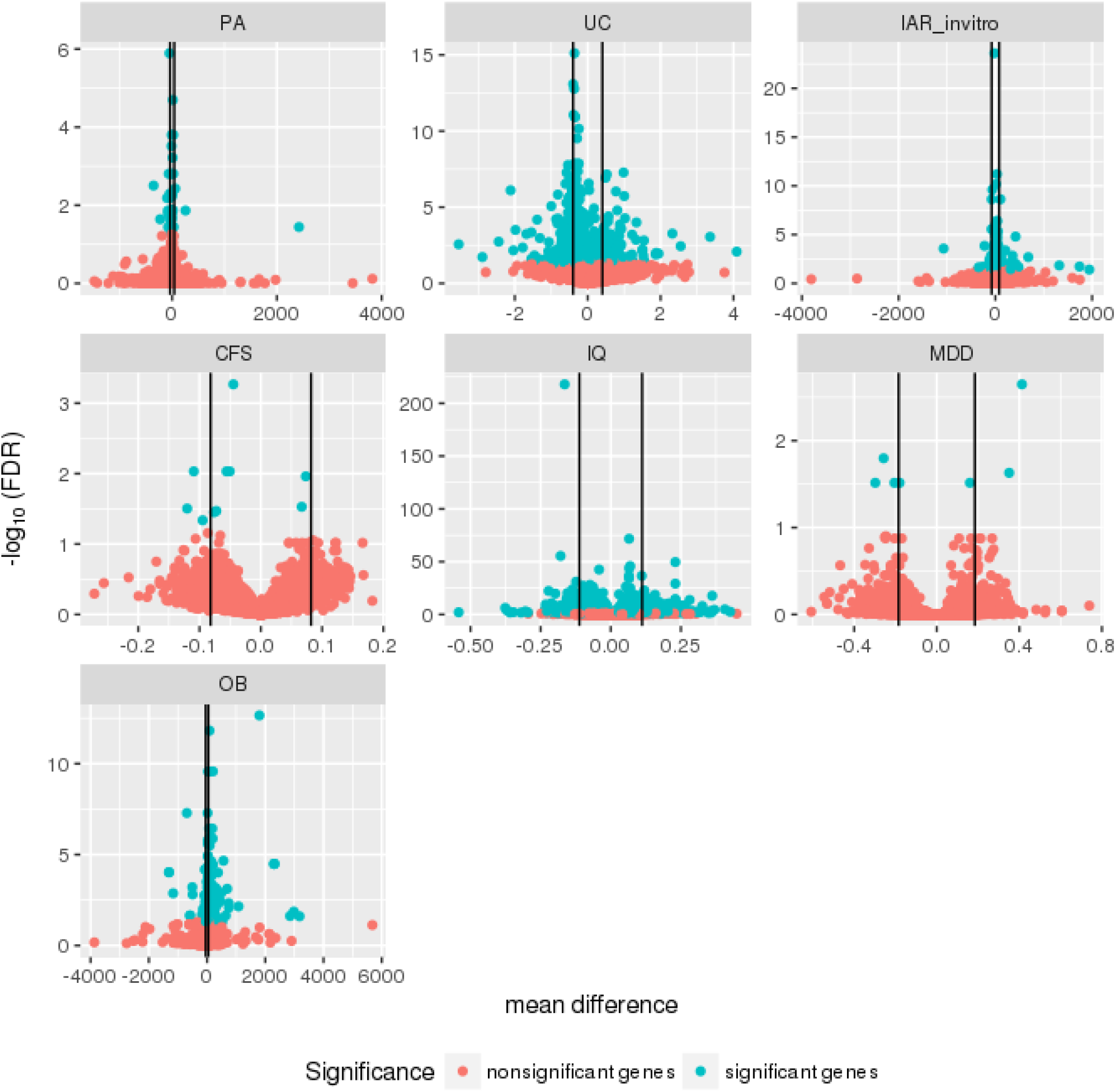
Volcano plots for seven phenotypes. The mean differences versus the negative log (base 10) of FDR for the seven phenotypes (each with greater than 10 twin pairs). The blue color indicates FDR significant genes (FDR < 0.05) and the red color indicates FDR nonsignificant genes. The black lines indicate the effect size thresholds (95th percentile of absolute value of mean expression differences for each phenotype).

Across the seven studies (phenotypes) incorporated into our analyses, we found that intelligence quotient (IQ) had 30 of the most significant genes (with FDR less than 0.05 and mean difference greater than the absolute value effect size threshold of the 95th percentile). Out of the disease phenotypes incorporated into our study, intermittent allergic rhinitis (IAR in vitro) had the most significant gene (*COQ5* [Coenzyme Q5, Methyltransferase], a gene involved in methyltransferase activity) with a FDR value of 2.4E-09 (mean difference = 105 units, or affected twins had higher gene expression than their unaffected twin pair).

The disease with the highest total number of significant genes out of the ones included in our analysis was UC (424 significant genes) and the one with the least was CFS (three significant genes). The non-disease phenotype with the highest total number of significant genes was IQ (677 significant genes) and the one with the least was physical activity (PA, 15 significant genes).

### Little overlap of differentially expressed genes in discordant twins across seven phenotypes

Next, we computed the pairwise similarity of gene expression between phenotypes in two ways. First, we computed the intersection between genes found significant between phenotypes. Second, we correlated the expression differences using a nonparametric Spearman correlation.

We report the percentage of the number of overlaps of significant genes out of the number of overlaps of measured genes for pairs of phenotypes (Table 2, S1, and S2). We found that the phenotype pair with the highest number of overlapping significant genes was UC and IQ (16 genes or ∼0.09% of total possible genes that overlapped, Table 2). The disease-disease pair with the most number of overlapping significant genes was OB and UC (13 genes or 0.06% of the total possible genes). The average percent of overlapping genes between phenotypes was 0.009%.

**Table 2.**
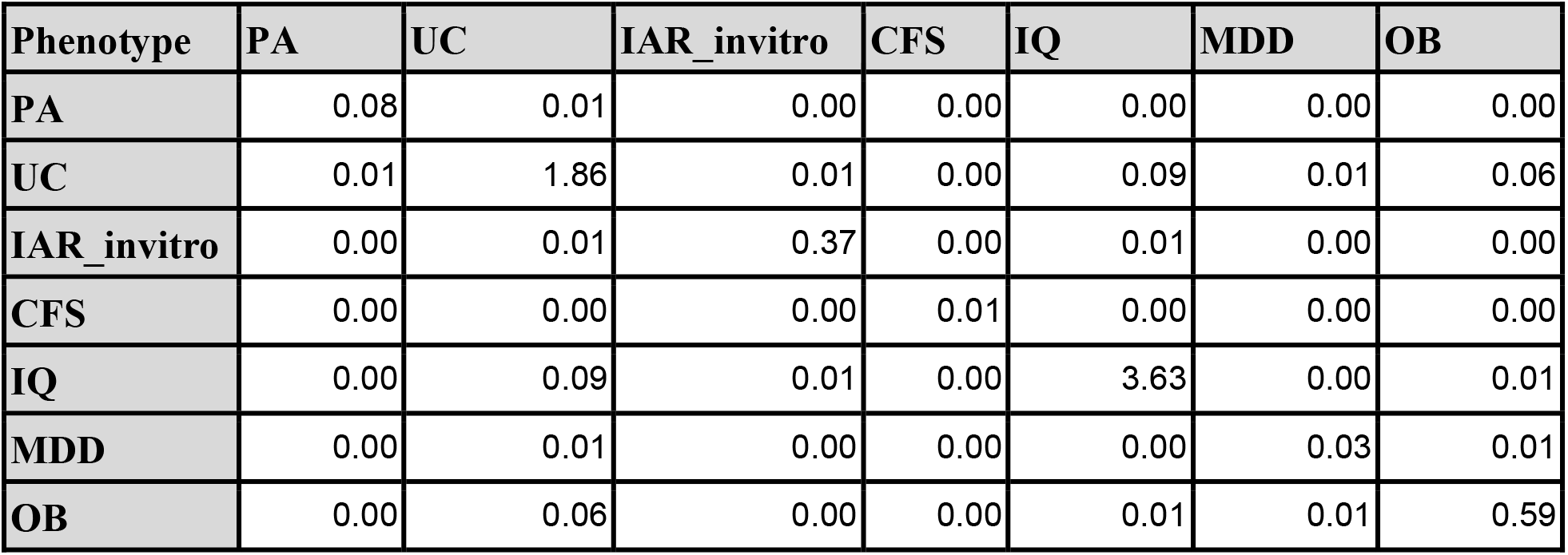
Percentages of Overlaps of Significant (FDR < 0.05 and Absolute Value Effect Size Threshold of 95th percentile) Genes. This table shows the percentages of overlapping significant genes in phenotype pairs out of the total overlapping measured genes in those pairs.

The pairwise Spearman correlations between the mean expression differences for each of the phenotype pairs were modest (Table 3). The absolute value of Spearman correlation coefficients ranged from 7.9E-4 to 1.8E-1.We found no significant correlations (with an unadjusted p-value threshold of 0.05) between the mean gene expression differences.

**Table 3.**
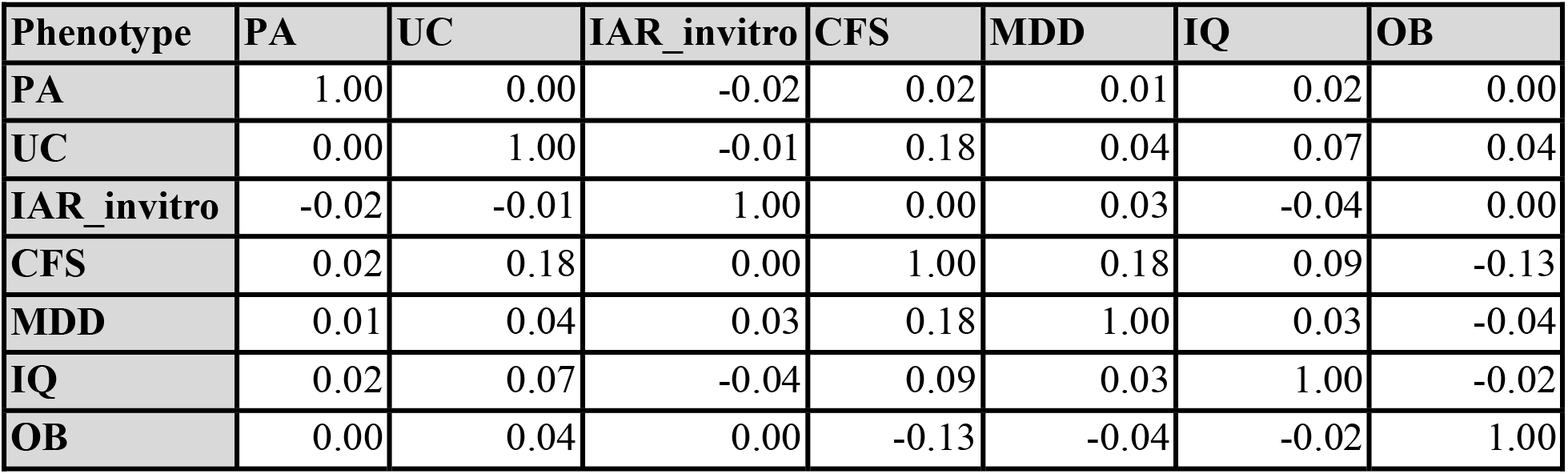
Spearman correlations of mean gene expression differences between phenotypes. This table shows the Spearman correlations between seven phenotypes in each phenotype pair.

| Phenotype | PA | UC | IAR_invitro | CFS | MDD | IQ | OB |
| --- | --- | --- | --- | --- | --- | --- | --- |
| PA | 1.00 | 0.00 | -0.02 | 0.02 | 0.01 | 0.02 | 0.00 |
| UC | 0.00 | 1.00 | -0.01 | 0.18 | 0.04 | 0.07 | 0.04 |
| IAR_invitro | -0.02 | -0.01 | 1.00 | 0.00 | 0.03 | -0.04 | 0.00 |
| CFS | 0.02 | 0.18 | 0.00 | 1.00 | 0.18 | 0.09 | -0.13 |
| MDD | 0.01 | 0.04 | 0.03 | 0.18 | 1.00 | 0.03 | -0.04 |
| IQ | 0.02 | 0.07 | -0.04 | 0.09 | 0.03 | 1.00 | -0.02 |
| OB | 0.00 | 0.04 | 0.00 | -0.13 | -0.04 | -0.02 | 1.00 |

### Discordant twin gene expression is heterogeneous across seven phenotypes

We hypothesized that it is possible to identify shared environmental etiology between phenotypes by identifying genes across multiple phenotypes. We performed meta-analysis (using the Dersimonian and Laird meta-analytic technique^8^) on each gene across all seven possible phenotypes to (1) estimate the overall mean difference of each gene across seven phenotypes (genes putatively expressed in greater than one phenotype) and (2) estimate how each gene’s mean expression difference varied across all of the seven studies (gene expression heterogeneity). The empirical cumulative distribution of meta-analyzed mean differences is shown in Figure S2. The I^2^ (heterogeneity) estimates versus the negative log (base 10) of FDR-corrected QEp values (measure of significance of I^2^ different from 0) is depicted in Fig. S3 and the empirical cumulative distribution plot of I^2^ values is shown in Fig. 3.

**Figure 3.**
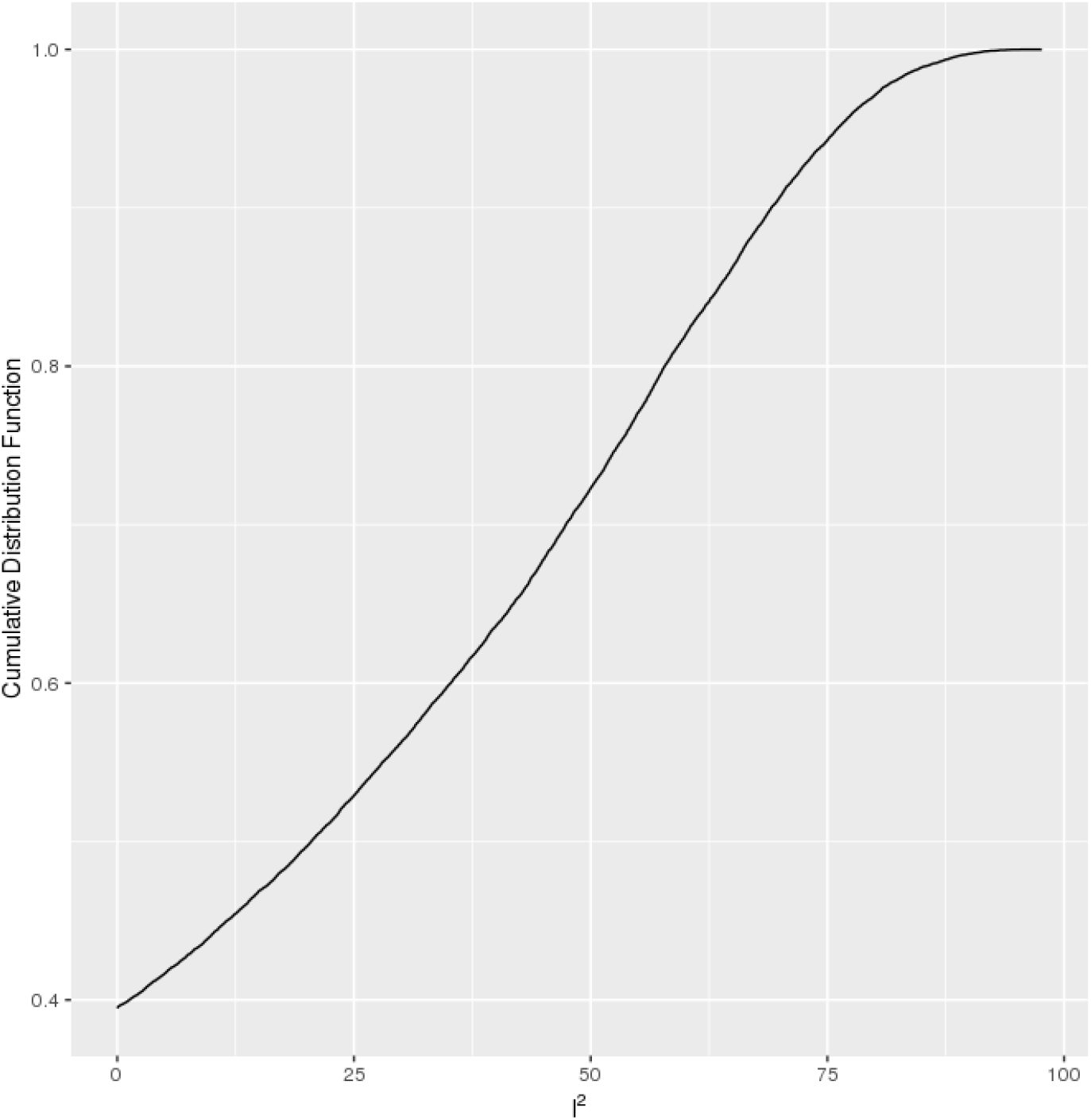
Empirical Cumulative Distribution Function Plot of I^2^ values. The distribution of all measured genes (from the seven studies) among their I^2^ values.

First, we discuss genes that were expressed over all phenotypes in discordant twins. We identified 19 out of the 25,154 total genes (0.08%) that were differentially expressed in discordant twin samples across multiple phenotypes (FDR-corrected p-value of mean difference less than 0.05, mean difference greater than the absolute value effect size threshold of the 95th percentile, and measured in more than one study; Fig. S2). The top significant differentially expressed genes (significant genes that were measured for multiple phenotypes) included those that are involved in keratinization such as *KRTAP19-5* (Keratin Associated Protein 19-5) and *KRTAP20-2* (Keratin Associated Protein 20-2). A third included *FGF6* (Fibroblast Growth Factor 6), a gene involved in normal muscle regeneration, all with FDR values less than 3.7E-4. We found no genes that were significant overall (FDR-corrected p-value of mean difference < 0.05, mean difference greater than the absolute value effect size threshold of the 95th percentile, and measured in more than one study) and that were also significant in individual disease phenotypes.

Second, we discuss overall heterogeneity of the differentially expressed genes. Out of all the 25,154 genes measured, 2,401 genes (10%) were found to have FDR-corrected QEp (measure of significance of I^2^) values less than 0.05, corresponding with I^2^ values of greater than 68%. None of the overall significant (FDR-corrected p-value of mean difference less than 0.05, mean difference greater than the absolute value effect size threshold of the 95 th percentile, and measured in more than one study) genes were found to also have FDR-corrected QEp values less than 0.05. Also,we found 40% of all measured genes to have I^2^ values of 0. In fact, 11 out of the 19 significant genes were found to have an I^2^ values equal to 0. The gene with the highest I^2^ estimate (47%) was *ZNF12*[Zinc Finger Protein 12] (mean difference of −0.13 and FDR-corrected p-value of mean difference of 0.04), a gene involved in transcription factor activity. We have little data to support that differential expression for most genes across multiple phenotypes is heterogeneous.

### Sex-specific gene expression heterogeneity across three phenotypes (MDD, OB, CFS)

We hypothesized that sex may play a role in differences in gene expression across twins. For the three phenotypes (MDD, CFS, and OB [Table 1]) that had samples labelled with sex, we carried out sex-specific differential expression analyses and meta-analyzed over each gene’s expression values (for each sex group separately).

For males, we found 9 overall significant genes (overall FDR-corrected p-values of mean difference < 0.05, mean difference greater than the effect size threshold of the 95th percentile of absolute value of mean expression differences, and measured in more than 1 phenotype). The most significant genes (with overall FDR values < 1.4E-7) were *PTPRN* (Protein Tyrosine Phosphatase, Receptor Type N), a gene involved in phosphatase activity and *TRNT1* (TRNA Nucleotidyl Transferase 1), a gene involved in nucleotidyltransferase activity. Next, we identified significant genes (using FDR and effect size thresholds) that had extremely low I^2^ (heterogeneity) estimates (0%) across male groups from all three phenotypes. We found all 9 genes to have I^2^ values of 0. Of those genes, the ones with the lowest FDR values were again *PTPRN* and *TRNT1*. All of the overall significant genes for males had I^2^ estimates equal to 0 %, suggesting that the significant genes in males have expression levels that are similar across different phenotypes.

For females, we found 12 overall significant genes (overall FDR < 0.05, mean difference greater than the absolute value effect size threshold of the 95th percentile, and measured in more than 1 phenotype). The most significant genes (with overall FDR values < 7E-3) were *RPL22* (Ribosomal Protein L22), a gene involved in poly(A) RNA binding, *WFDC1* (WAP Four-Disulfide Core Domain 1), a gene involved in growth inhibitory activity, and *DHX40* (DEAH-Box Helicase 40), a gene involved in helicase activity. Out of the 12 overall significant genes, we found 11 genes to have I^2^ of 0. The overall significant gene with the highest I^2^ estimate (99%) was *MS4A4A* (Membrane Spanning 4-Domains A4A). Except for *MS4A4A*, we found that all other female-specific overall significant genes had I^2^ estimates equal to 0%. While we found similar number of genes differentially expressed in male and female groups of discordant twin pairs, there was no overlap between the sexes.

### Checking for Batch Effects

We were cautious of possible batch effects impacting our analyses. In order to check for possible batch effects, we utilized the COmbat CO-Normalization Using conTrols (COCONUT^29^) tool to batch correct the samples and ran our differential analysis pipeline on these batch-corrected samples. We compared the results of the samples prior to the correction with the results after the correction by correlating the mean differences obtained before and after correction. The correlations ranged from 0.72 (OB) to 0.95(IQ) (Table S3). We also specifically computed the Spearman correlations between the mean differences obtained before and after correction of genes found significant (only using FDR < 0.05 threshold in differential analyses prior to correction). These correlations ranged from 0.88(UC) to 1(MDD) (Table S4). We did not have evidence to support that batch correction would significantly alter the findings. Therefore, to increase sensitivity of the number of genes queried, we decided to report non-corrected results.

## Discussion

We present a computational workflow to execute differential expression and heritability analyses using discordant twin samples across multiple (previously) disparate phenotypes. In addition, our work has resulted in an online database resource (*DiscTwinExprDB :* http://apps.chiragjpgroup.org/disctwinexprdb/) for researchers to query differentially expressed genes in discordant twins. Briefly, we first identified genes differentially expressed in seven different phenotypes (each with at least ten twin pair samples) by finding differences in transcript-level (probe-level) expression values and meta-analyzing those transcript-level differences to get overall gene-level differences using monozygotic (MZ) discordant twin samples.

There have been multiple investigations (whose data have been deposited in the GEO and dbGaP repositories) that have been published recently utilizing the MZ discordant twin study design to perform differential expression analyses such as Byrnes *et al*^9^ (GEO: GSE16059). It is important to note that most of these studies have reported transcript-level (probe-level) values. For most microarray platforms, multiple probes map to the same gene and each probe sequence has different binding affinity leading to ‘different measurement scales’ (Ramasamy *et al*.^10^). Hence, rather than reporting the transcript-level values, summarizing those values to gene-level values by meta-analyzing over each gene’s corresponding probe-level values may yield more interpretable results^10^. While studying expression at the transcript or gene-level is of debate^10^, here, rather than studying differential expression on the transcript level, we used the meta-analytic gene-level summarization technique (as has been implemented previously by Sweeney *et al*.^7^). We hypothesize that the use of this technique provides better power than alternative transcript-level methods; in fact, we showed that we were able to detect more genes (found total of 10 significant genes) than the Byrnes *et al*^9^ investigation (this study detected none).While we claim investigating gene-level expression differences enhances interpretability, further study must be devoted to systematically test what methods yield more power.

We note the variation in the magnitude of mean expression differences across the different phenotypes and studies. The inter-study variation may be explained by the diverse platforms, sample preparation and processing, sample types, and sample sizes of the different studies. Furthermore, the variation may also be explained by the vast potential differences in mechanisms of environmental exposure or phenotype itself on gene expression. To enhance comparability across studies, we also reported the ranks of the gene expression differences within each study in our R Shiny web application.

Our study is also the first -- to the best of our knowledge -- to carry out differential gene expression analysis in MDD discordant twin pair samples. We identified five significant differentially expressed genes. Many of these genes have not been identified as being directly associated with MDD in scientific literature before and these genes need to be replicated. However, a few of these genes were mentioned as being associated with depression and alternative splicing of exons (for example *TRA2B*^11^).

One factor that may play a role in population-level differences in gene expression includes sex. By stratifying our analyses by sex, we have identified multiple significant differentially expressed genes in the sex-specific analyses that appear to be involved in protein and RNA binding activity. For example, *PTPRN* and *TRNT1*, which we have identified to be significant in the male-specific analyses, both appear to play a role in enzyme binding activity. We also observed that in the female-specific analyses, the most significant genes appeared to be enriched for RNA binding. There was little to none overlap of overall significant genes between the sexes. We found an intersection of 2 genes between the gender groups with a less stringent effect size threshold initially and after using a more stringent absolute value effect size threshold (95th percentile of absolute value of mean expression differences) we found no overlap. It is unclear why such a difference manifests between sexes and more investigations must commence to uncover the causative role of sex in phenotype^12^.

From our overall meta-analysis, we identified genes across various studies with low I^2^ (heterogeneity) estimates and relatively few with higher I^2^. For example, we found that genes such as those involved in keratinization were expressed across all of the phenotypes (e.g. *KRTAP19-5*), but we lacked power to implicate these genes in any single phenotype. Second, these genes exhibited little heterogeneity and were not found significant in any one phenotype, suggesting a critical role across phenotypes. As the reader may know, these genes are known to play an integral role in hair shaft formation^13^. Most of the overall significant genes were found to have little heterogeneity (variance across multiple phenotypes) suggesting that, while there may be a common discordant differential gene expression signature across disparate phenotypes, their effect sizes (difference) is modest.

In conclusion, we found genes significantly expressed in discordant twins are specific to phenotype and we have little evidence to support shared environmental etiology between these seven phenotypes. For example, we found little overlap between genes expressed in disparate phenotypes (on average, 0.009% of co-measured genes). Second, from our meta-analysis, we identified overall common differential gene expression signatures for the phenotypes, but these signatures were not identified in individual phenotypes. This suggests that if there is a common thread of gene expression across disparate phenotypes, the effects are probably small.

Our work has multiple limitations. One limitation to our and other studies using monozygotic discordant twin pair samples is that they are relatively small in number (sample size range for individual phenotypes was 10 to 44 pairs and total sample size was 92 pairs). It is a challenge to recruit and assay identical twins and even collect publicly available data for a given phenotype. Relatedly, and critically, genes we report need to be followed up and replicated in other twin investigations. Our second limitation was the lack of adequate datasets with sufficient sex information to untangle the role of sex in differential gene expression. Third, while studying MZ twins is a natural way to control for the role of inherited factors in gene expression variation, we cannot rule out the role of the disease or phenotype itself in modulating gene expression (e.g., reverse causality). One phenotype for which this is readily apparent is physical activity. For example, it is possible that the phenotypic state itself may induce changes in gene expression decoupled from environmental influence. A way to mitigate the chance of reverse causation for disease-related phenotypes includes incorporating time into the analysis, such as following twin pairs through the life course. In the future, we aim to collect twin data in an unbiased manner to ascertain the role of reverse causation in expression to deconvolve the role of phenotype on gene expression change. We emphasize that more resources devoted to the functional and biological differences between discordant and concordant twins should be developed and made available to enhance replication and study design, such as the impactful TwinsUK cohort^14^. To enable investigations across the studies analyzed here, we provide a web-accessible database (*DiscTwinExprDB*) for straightforward reuse of our analyses in other integrative contexts.

We hope that our work will inspire future studies to further understand the role of the environment in multiple phenotypes, eventually leading to the identification of environment-specific influences in multiple disease phenotypes.

## Methods and Materials

A schematic diagram depicting our analysis workflow is shown in Fig. 1.

### MZ Discordant Twins’ Gene Expression Data

We collected gene expression datasets from the Gene Expression Omnibus^4^ (GEO : https://www.ncbi.nlm.nih.gov/geo/) [Table 1]. The other data sources we used for our analyses were the Database of Genotypes and Phenotypes^5^ (dbGAP) and Array Express^6^ (AE). We used the phs000486.v1.p1 (“Integration of Genomics and Transcriptomics in unselected Twins and in Major Depression”) study from dbGAP and the E-MEXP-1425 study from AE.

Our data selection process is depicted in Fig. 1. We selected 5 monozygotic discordant twin studies (measuring discordance for phenotype) with at least 10 pairs in each of the studies from GEO (Table 1), which provided 92 MZ discordant twin pairs to analyze the gene expression samples from. We filtered out studies that had ambiguity in reporting (e.g., for some samples it was unclear as to which samples constitute a twin pair and what phenotypic status the samples had) and those that had with very low sample sizes (number of twin pairs < 10). We filtered out 29 such studies.

We downloaded one study (phs000486.v1.p1) from dbGAP that has 28 MZ discordant twin pairs for major depression. Further, we downloaded a dataset (E-MEXP-1425) from ArrayExpress (AE) on Obesity that had 13 MZ discordant twin pairs. The summary of all of these datasets that were used are shown in Table 1.

For the sex-specific analyses, we used 1 GEO study (GSE16059, CFS) along with the AE study (E-MEXP-1425, OB) and dbGAP study(phs000486.v1.p1, MDD), as these were the only three studies that had systematically provided the corresponding gender attribute information.

We wrote R scripts to download and transform these expression datasets into a compatible format for our analyses (https://github.com/stejat98/disctwinexpr/).

### Phenotype-specific analysis of genes differentially expressed in twins

In summary, we performed differential gene expression analysis on the seven different MZ discordant twin gene expression studies across a total of 25,154 genes and identified a list of significant genes (using a false discovery rate [FDR] threshold of less than 0.05 and effect size threshold of 95th percentile [of absolute value of mean expression differences in each phenotype]) for the phenotype being observed in each study.

Specifically, we executed a paired t-test on the twins’ gene expression (probe- or transcript-level) data in each of the seven monozygotic discordant twin gene expression studies separately. We obtained the mean differences, the standard errors, and the p-values for each microarray probe. Next, we mapped each probe to its corresponding gene using the annotation tables for each of the platform types used by the seven different studies. We performed a fixed-effect meta-analysis across probe-level values for each gene (inspired by Sweeney *et al*.^7^) using the R ‘rmeta’ package^28^. This yielded the overall mean differences, standard errors, and the p-values for each gene for each of the seven studies. Since the seven studies used different microarray platforms, the numbers of total measured genes were different from study to study (the number of pairwise measured genes is in Table S2). To enhance comparison among the seven studies, we also computed the rank order of the expression differences in each study (available in the R Shiny web application: http://apps.chiragjpgroup.org/disctwinexprdb/).).

Last, we also performed FDR (False Discovery Rate [Benjamini-Hochberg]^15^) correction on the p-values for each gene for each study. We identified significant genes within each study using a FDR threshold of 0.05 and effect size threshold of the 95th percentile (of absolute value of mean expression differences in each phenotype [Table S5 for thresholds]) and deemed these as “significant” in our study. We leveraged the same pipeline to identify sex-specific expression differences. Specifically, we carried out the paired t-test and the meta-analytic gene-level summarization for each sex group separately, using the CFS, OB, and MDD datasets.

We performed meta-analysis on each gene-specific value across all studies to measure how each gene’s mean expression difference varied across the seven studies for each of the 25,154 unique genes measured in total (across all studies). This was done by using the Dersimonian and Laird meta-analytic technique with the *‘metafor’* package^16^ (rma.uni function).We also produced forest plots to illustrate the variance of expression levels for each gene (http://apps.chiragjpgroup.org/disctwinexprdb/). The meta-analysis yields two important metrics for measuring heterogeneity (I^2^ and QEp).The I^2^ estimate is a commonly used metric to measure percentage of variation in meta-analysis that is due to the actual heterogeneity of the studies included in the meta-analysis^17^. The QEp is a statistic (p-value) used to measure the significance of heterogeneity (with the null hypothesis that there is no heterogeneity) ^18^.

## Code availability

We made our code accessible at (https://github.com/stejat98/disctwinexpr/).

## Web Application

We built a web application to visualize the results from our analyses using the Shiny Web Application Framework in R that is accessible at (http://apps.chiragjpgroup.org/disctwinexprdb/).

## Acknowledgements

C.J.P. and S.T. are funded by NIH grants (ES023504 and ES025052). C.J.P. is also funded in part through a NSF Big Data Spokes grant (NSF 1626870).

## Author Contributions

S.T. carried out the analyses. C.J.P. served as mentor and advisor for the project. S.T. and C.J.P wrote the paper.

## Additional Information

### Competing financial interests

The authors report no conflicts of interest.

